# No evidence of *Toxoplasma gondii* infection in urban and rural squirrels

**DOI:** 10.1101/2020.09.28.313866

**Authors:** Riikka P Kinnunen, Chloé Schmidt, Adrián Hernández-Ortiz, Colin J Garroway

**Author notes:** Correspondence*, Riikka Kinnunen, Colin Garroway.

## Abstract

Wildlife in cities can act as reservoirs for parasites that can infect domestic pets and have implications for human health. For example, the Coccidian parasite *Toxoplasma gondii* can infect all mammalian species, with felids acting as the only definitive hosts. Rodents, such as squirrels, could play an important role in urban infection dynamics of *T. gondii*, as squirrels occur at higher densities in cities than in natural environments and regularly share their territories with cats. In urban and suburban areas squirrels can encounter infectious oocysts shed in cat feces in contaminated soil or in the food they eat. They are also preyed upon by cats making them a potentially important species for completing the *T. gondii* life cycle in cities. We hypothesized that due to increased exposure to cats, urban squirrels would be more susceptible to *T. gondii* infection relative to squirrels in more natural areas. We investigated this using molecular and serological methods on samples collected from American red squirrels (*Tamiasciurus hudsonicus*), eastern grey squirrels (*Sciurus carolinensis*), northern flying squirrels (*Glaucomys sabrinus*), and least chipmunks (*Tamias minimus*) in and around the city of Winnipeg, Manitoba, Canada. We tested a total of 230 tissue samples from 46 squirrels for *T. gondii* DNA using quantitative PCR and supplemented these data with analyses of 13 serum samples from grey squirrels (*Sciurus carolinensis*) testing for *T. gondii* antibodies by indirect ELISA. We found no evidence of *T. gondii* infection in any squirrel. This suggests that squirrels may not be important intermediate hosts of *T. gondii* in cities and do not need to be considered as sources of infection to cats.

## Introduction

Wildlife in cities can act as reservoirs for parasites that can infect domestic pets and have implications for human health [1]. Indeed, previous work has found increased levels of wildlife parasitism in cities compared to rural areas [2–5]. This increased level of parasitism could reflect both the increased population density of urban host species and higher within and between species contact rates in response to resource provisioning in cities relative to rural areas [6,7]. Contrasting this pattern, in some cases the overall species richness and diversity of parasites can be reduced in cities as urbanization can lead to a decline in host species richness. With reduced access to hosts, parasites with one or a few host species may then also become extirpated [7]. Parasites that spread through direct contact or oral-fecal routes are likely to be favored in urban areas, [7] but general knowledge underlying the ways host-pathogen interactions operate in cities is still limited [1].

The Coccidian parasite *Toxoplasma gondii* is an interesting parasite within the context of urban ecosystems. Domestic cats—numerous in cities as pets [8,9]—and other Felidae are the only known definitive hosts of *T. gondii* [10]. When infected, cats can shed millions of infectious *T. gondii* oocysts into the environment daily in their feces for a duration of one to two weeks, making them an important part of the parasite’s life cycle [11,12]. Humans can acquire infection from cats by accidentally ingesting oocysts, for example when cleaning cat litter or not washing hands after gardening [13]. Other mammals can act as intermediate hosts for *T. gondii*, acting as reservoirs for the parasite and, in some cases, as possible sources of infection for cats in cities. Cats and intermediate hosts can become infected by consuming carcasses infected by *T. gondii* tissue cysts, or by coming into contact with oocyst contaminated soil, water, or food. A significant proportion of the human population globally is infected with *T. gondii*, but most healthy people do not experience symptoms of infection [14]. However, immunocompromised people and pregnant women require medical intervention to avoid serious health issues [14–16]. Consequently, further knowledge of *T. gondii* infection dynamics in cities is needed.

Urbanization can increase the risk of an animal being exposed to *T. gondii* [4,17,18]. Many squirrel species (Sciuridae) are ubiquitous in cities, where they are commonly found at much higher densities than in natural environments [19]. In urban and suburban areas squirrels regularly share their territories with domestic cats and collect and cache their food in backyards and gardens where they can encounter infectious oocysts shed in cat feces in contaminated soil or in the food they eat. This may make urban squirrels particularly susceptible to parasite infection compared to their rural counterparts. After being infected squirrels act as intermediate hosts for the parasite, and the parasite can remain within the host body in tissue cysts for the rest of the host’s life. The infection can be asymptomatic or develop into the disease toxoplasmosis [20,21]. Many *T. gondii* strains isolated from nature are of low virulence, leading to subclinical toxoplasmosis that does not kill the animal, but can make prey—such as a squirrel—susceptible to predation by cats [22], thus enabling the parasite to complete its life cycle [20,22]. Squirrels may thus act as a source of infection to cats, in a similar way to other prey species [23]. As such squirrels may play a role in *T. gondii* population and infection dynamics in cities.

Our aim in this paper was to survey the prevalence of *T. gondii* infection in squirrel (Sciuridae) populations in and around the city of Winnipeg, Manitoba, Canada. We specifically asked whether squirrel species (American red squirrel, *Tamiasciurus hudsonicus*; eastern grey squirrel, *Sciurus carolinensis*; northern flying squirrel, *Glaucomys sabrinus*; and least chipmunk, *Tamias minimus*) are important intermediate hosts of *T. gondii* and whether *T. gondii* infection is more common in a city than in more natural habitats. We hypothesized that, due to high population densities of both squirrels and cats in cities [9,19], urban squirrels may act as intermediate hosts of *T. gondii* and that urban squirrels will have a higher prevalence of *T. gondii* infection than rural squirrels. We tested these hypotheses by using molecular and serological methods on samples collected from four squirrel species in and around the city of Winnipeg, Manitoba, Canada.

## Materials and Methods

To investigate *T. gondii* prevalence in squirrels in chronic and acute stages we followed recommendations from previous studies and used complementary molecular and serological methods [24]. Quantitative PCR was done on a wide-ranging sample from four squirrel species (American red squirrel; eastern grey squirrel; northern flying squirrel; and least chipmunk) and serological testing on focal study populations of grey squirrels in and around the city of Winnipeg, Manitoba, Canada (Fig. 1). Winnipeg is the largest city in the province of Manitoba with a population of 778,489 and a total land area of 464,33 km^2^ [25]. Winnipeg lies 239 meters above sea level and has high seasonal climatic variation, with temperature varying from the extremes ranging between −24 °C to −33 °C between January to March to around +30 °C to +35 °C between June to September [26].

**Figure 1.**
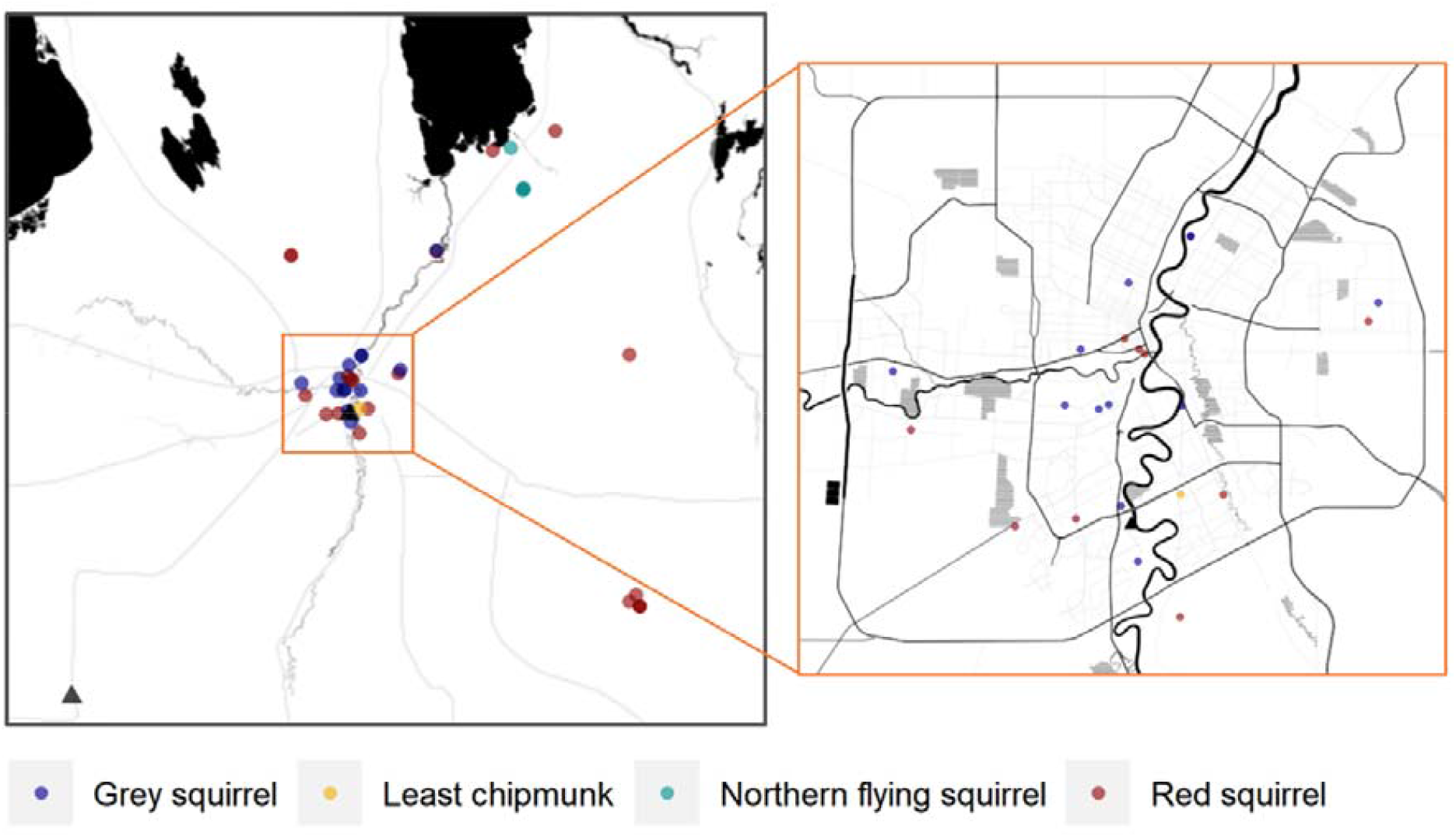
Map showing the sampling locations of squirrel (Sciuridae) carcasses (circles) used to collect tissue samples for quantitative PCR detection of *Toxoplasma gondii*, and the urban and rural study sites for blood collection (triangles) from grey squirrels (*Sciurus carolinensis*) for serological testing of *Toxoplasma gondii* antibodies. The map on the left shows the urban and rural locations of squirrels from in and around the city of Winnipeg, Manitoba, Canada, and the map on the right is a close-up of the urban locations within the city perimeter. The map was created using an R package *ggmap* [27]. Map tiles by Stamen Design, under CC BY 3.0. Data by OpenStreetMap, under ODbL.

### Collection of Carcass Samples

To investigate *T. gondii* prevalence in squirrels during the chronic stage we collected the carcasses of 25 red squirrels, 16 grey squirrels, four flying squirrels, and one chipmunk from trappers, wildlife rehabilitation centers, and pest control companies during the years 2017 to 2019 to collect tissue samples for PCR (Fig. 1). We did not obtain the carcasses close enough to death to collect usable serological samples.

### Molecular Methods

Upon necropsy, we collected the entire liver, spleen, brain, heart, kidneys, and lungs, and stored tissues separately at −20 °C until further analysis. As *T. gondii* is a cyst-forming parasite, the detection probability of *T. gondii* can differ between organs [28]. Consequently, we tested multiple samples per individual from two to six different organs to maximize the probability of detecting the parasite.

#### Cell lysis

Cell lysis was done by first adding 3 ball bearings to each 2 mL screw-cap tube containing 0.6 mL of ATL buffer, and adding 100 mg of frozen tissue or pipetting 0.1 mL of sample (if liquid such as a thawed brain) to the tube. We placed the samples in a BeadBeater for 3 minutes after which they were quickly centrifuged. We added 70 µL of Proteinase K to the ATL lysate. We incubated the lysate at +56 °C for 1-3 hours during which the tubes were intermittently inverted several times. We centrifuged the lysate quickly and added 0.6 mL of AL buffer. We inverted the tubes several times and incubated them at +70 °C for 10-30 minutes in a dry block. We then again inverted the tubes intermittently several times. The samples were then centrifuged for 3 minutes at 10,000 x *g*. After the completion of the cell lysis, we used 100 µL of the ATL/ProtK/AL lysate to continue the extraction.

#### Nucleic acid extraction and real-time PCR

We did nucleic acid extraction for *T. gondii* using 5X MagMAX viral Isolation kit (Applied Biosystems AMB1836-5). Primers and probes for real-time PCR were designed according to the protocol of De Craeye et al. (2011) [29] (see Table S1 for primers and probes). PCR product size was 106 bp. We used cellular r18S (ribosomal RNA gene) as an internal control of all PCRs. Real-time PCR was conducted using TaqMan Fast Advanced Master Mix (Applied Biosystem) in Applied Biosystems™ 7500 Real-Time PCR System. Each PCR reaction had concentrations of 10 µM of T2 /F primer, 10 µM of T3/R primer, and 5 µM of probe in the master mix. Thermo cycling Program: Initial denaturation and activation of the Taq polymerase at +95 °C for 2 minutes, followed by 45 cycles at +95 °C for 5 seconds and +60 °C for 33 seconds. We analyzed the results using 7500 System SDS Software.

### Serum collection

To investigate *T. gondii* prevalence in squirrels during the acute stage we conducted live trapping of grey squirrels in one urban and one rural site between 5^th^ June and 1^st^ August 2019 (Fig. 1). The urban site is located in the city of Winnipeg, Manitoba, Canada, and consists of an ∼10 ha park located on the University of Manitoba campus and a suburban neighborhood next to the park. The study site is bordered by the Red River and two major highways with high amounts of car traffic. The rural site is a ∼34 ha forest patch next to an active honey-farm, near the twin cities Morden and Winkler in southern Manitoba (49°24’01.1”N, 98°00’29.2”W), bordered by agricultural land. We used live traps (Tomahawk Live Trap Co., Tomahawk, WI, USA) to capture grey squirrels at the study sites. Between 20-40 traps in total were set to the urban site and between 80-100 traps to the rural site each trapping day. Traps were baited with peanut butter and checked regularly. The number of traps was different for the two sites to ensure approximately the same number of squirrels was captured at both sites. Traps were placed in sheltered locations under vegetation cover or covered with canvas to provide shade and to calm the animals when in the trap. After capture, squirrels were handled in a canvas capture bag and we recorded the weight (g), body and tail length (cm), skull width (cm), age (adult or juvenile), reproductive status, and sex for each individual. We collected a minimum of 500 µL of blood from the femoral vein of each grey squirrel and stored the sample on ice until processing. Each squirrel was pit-tagged between the shoulder blades with passive integrated transponder (PIT) tags. All efforts were made to minimize suffering. We then released the squirrels at the place of capture. Our protocol (Protocol Number: f16-003) was approved by the University of Manitoba animal care and use committee following Canadian Council on Animal Care guidelines.

### Serological Methods

We collected a minimum of 500 µL of blood from 15 individual grey squirrels and stored samples on ice. All samples were processed within 12 hours. We centrifuged the collected blood samples at 3500 rpm for 15 minutes and froze the serum at −20 °C until used for testing.

#### Enzyme-linked immunosorbent assay

We used enzyme-linked immunosorbent assays (ELISA) to detect serum antibodies (IgG) against *T. gondii*. As species-specific conjugates are not available for squirrels, we used a commercially available ELISA kit for testing the samples (Multi-species ID Screen Toxoplasmosis Indirect kit, IDVet, Grabels, France) following the manufacturer’s instructions. We read the optical density values at 450□nm in a spectrophotometer and calculated results using these values and kit controls expressed as S/P (Sample to Positive Ratio) percentage (S/P%). We considered samples with S/P% less or equal to 40% negative; samples with S/P% between 40 and 50% doubtful or inconclusive; and samples with an S/P% higher than 50% positive, following the kit’s protocol.

## Results

We tested a total of 230 tissue samples from 46 squirrels from four squirrel species for *T. gondii* DNA using quantitative PCR (Table 1). Twenty-six of the carcasses were from urban locations within Winnipeg and 20 were from rural locations between 30 and 250 km from Winnipeg (Fig. 1). We had 25 American red squirrels (*Tamiasciurus hudsonicus*); 16 eastern grey squirrels (*Sciurus carolinensis*); four northern flying squirrels (*Glaucomys sabrinus*); and one least chipmunk (*Tamias minimus*). *T. gondii* DNA was not detected in any of the 230 tissue samples (liver; heart; brain; lung; spleen; kidney).

**Table 1.**
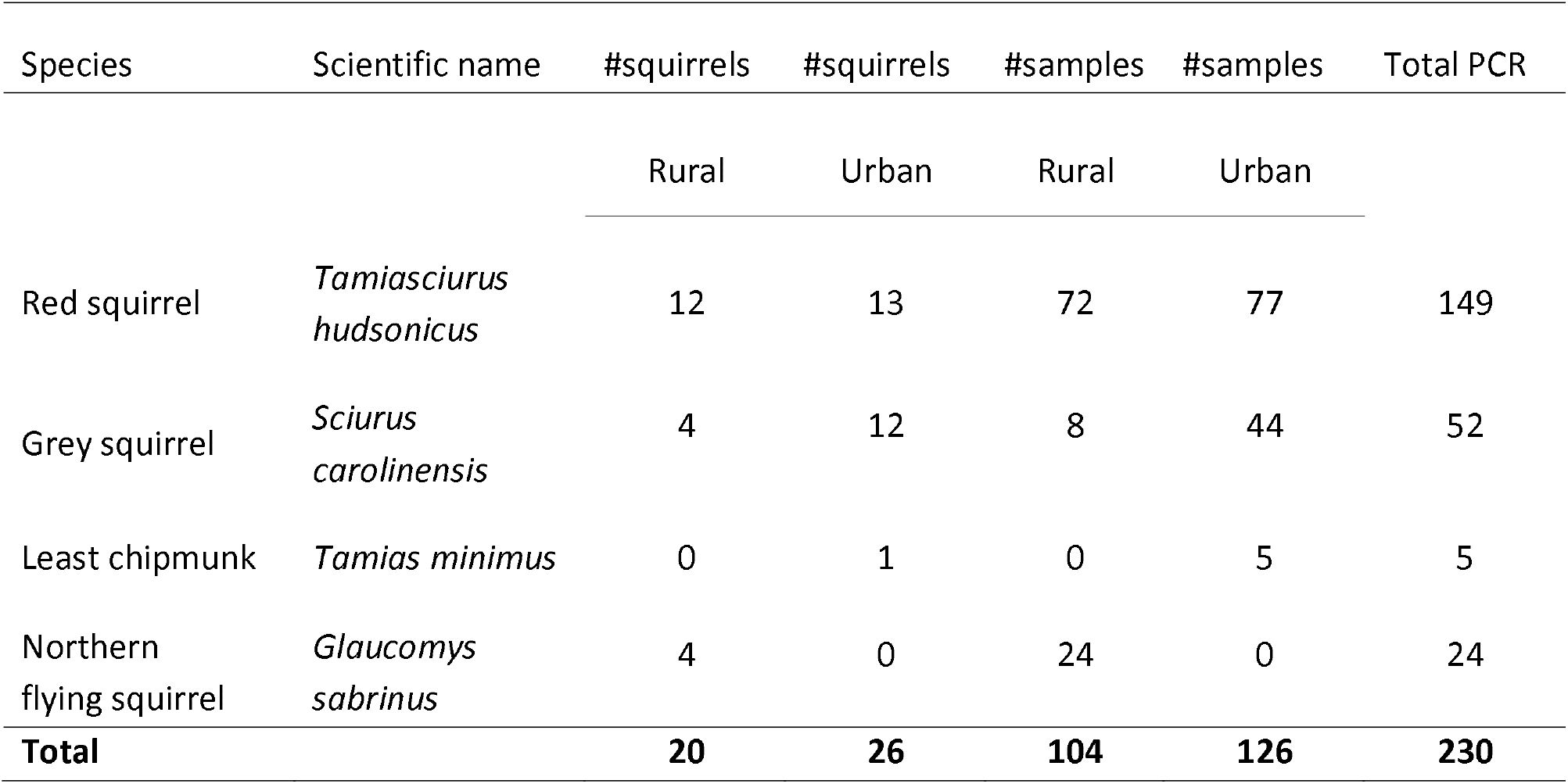
The number of individuals and the number of tissue samples from each squirrel species from rural and urban sampling locations tested for *Toxoplasma gondii* DNA using quantitative PCR. The common and scientific names of species are listed, as are the total numbers of PCR samples tested for each species, and the total number of individuals and samples in each sampling location.

We also tested a total of 13 (out of 15) samples of blood sera from grey squirrels for *T. gondii* antibodies (IgG) by indirect ELISA. Two of the collected samples did not have enough volume for testing. These results were negative—no *T. gondii* antibodies were detected in any sample (see Table S2).

## Discussion

We hypothesized that, due to higher population densities of both squirrels and cats in cities [9,19], urban squirrels may act as important intermediate hosts of *T. gondii* and have a higher prevalence of *T. gondii* infection than rural squirrels. However, we found no evidence of *T. gondii* infection in squirrels.

*T. gondii* has been found in many wild animals including on farms and in natural areas (e.g., gray fox (*Urocyon cinereoargenteus*) [30–32], red fox (*Vulpes vulpes*) [33]). However, studies investigating *T. gondii* infection dynamics in cities are still relatively rare [4,17, but see e.g., 18,34–37] considering the parasite can infect most mammalian species, including domestic cats and humans. Rodents can act as intermediate hosts for the parasite and as sources of infection to cats if consumed when infected by *T. gondii*. Squirrels are among the many species capable of getting a *T. gondii* infection, but our knowledge is limited, and studies have mostly focused on acute, fatal cases with little information existing on host-pathogen dynamics during chronic, latent infection [21].

The prevalence of *T. gondii* in Sciurids in urban areas is not well known. An earlier study from Guelph, Ontario, Canada found no evidence of infection across nine locations within and around the city from eastern grey squirrels (n= 16, the number of urban captures not specified) and chipmunks (*Tamias striatus*, n= 6) using the Sabin-Feldman dye test [30]. In natural areas, toxoplasmosis has been found in the eastern grey squirrel [20,38–40], western grey squirrel (*Sciurus griseus*) [41], and Eurasian red squirrel (*Sciurus vulgaris*) [12,21]. However, *T. gondii* prevalence in squirrels has generally been low (indirect hemagglutination test 1 of 265 positive individuals [42]; PCR 3 of 19 [21]; Sabin-Feldman dye test 2 of 11 [40]).

Using serological techniques with bioassay and PCR together can give a more reliable estimation of infection rate [24], as *T. gondii* is a cyst-forming parasite, and the distribution of *T. gondii* tissue cysts can be uneven and vary between organs [28]. This can lead to detection difficulties using PCR-based methods [43]. The sensitivity and specificity of antibody detection methods can also vary, which can lead to misinterpretation of results and possible false negatives and positives [44]. The multi-species ELISA kit we used has been successfully used to detect *T. gondii* antibodies in wildlife [45,46], has high sensitivity and specificity compared to other serological tests such as the modified agglutination test [47], and does not cross-react with other coccidian parasites—a factor known to limit the specificity of serological assays [48]. As we sampled several different organs and tissues per individual by PCR and used serological methods as an additional test to survey *T. gondii* prevalence in squirrels our results are less likely to be false negatives.

*T. gondii* prevalence in Manitoba, in general, is not well known and no previous survey of *T. gondii* prevalence in squirrels exists from the province. Serological testing in 1981 reported that of 55,527 pregnant women 129 showed signs of a recent *T. gondii* infection (Sekla et al. 1981). The same study also reported that 19 of 72 cats and one polar bear (*Ursus maritimus*) tested positive for *T. gondii* but results from 28 other species were all negative. An earlier study from Manitoba in 1976 found that *T. gondii* prevalence in pregnant women in urban areas was 8.17 %, and 6.29 % in rural areas [50]. The prevalence of *T. gondii* can be high in domestic sheep, pigs, and cattle [e.g., 30,51, however, see 52], yet studies from Manitoba or neighboring Saskatchewan have mostly found low prevalence [53,54].

The extreme climatic variations in the region may decrease the viability or infectivity of oocysts and tissue cysts in carcasses, therefore decreasing overall *T. gondii* prevalence in the area [53]. The temperature in Winnipeg between June to September can reach extremes of +30 °C to +35 °C [26] with daily average temperatures varying from approximately +20 °C to +13 °C [55]. *T. gondii* oocysts are highly resistant to environmental variation but sporulation (i.e. infectivity) is dependent on fixed temperatures and factors such as soil moisture that can influence the time oocysts can survive at high temperatures [56]. Additionally, winters in Winnipeg can be cold and windy with extreme temperatures ranging between −24 °C and −33 °C between January to March. Oocysts cannot sporulate and become infective after exposure to −21 °C for 1 day or −6 °C for 7 days [57]. After sporulation, oocysts can withstand lower temperatures better, being able to survive at −21 °C for 28 days [57], yet oocysts can not sporulate if the conditions are unfavorable [56]. It is thus possible that the combination of hot summers and cold winters reduces the viability of oocysts in Manitoba. Contact rates between domestic cats and *T. gondii* intermediate hosts may also be lower during the winter if people keep their pets indoors in cold weather. Winnipeg has an average yearly precipitation of 521 mm, but the years 2018 and 2019 were unusually dry in southern Manitoba [58]. This could have influenced oocyst survival in the area during data collection, as oocysts survive better in moist than in dry conditions [57,59].

We note that *T. gondii* antibodies have been found in skunks (*Mephitis mephitis*) and raccoons (*Procyon lotor*) from Saskatchewan (modified agglutination test: average seroprevalence in skunks 15.6%; in raccoons 20.8% in 1999, 12.5% in 2000) and Manitoba (in skunks 28%; in raccoons 27.5%) [60], which implies that although climatic variation may lower the prevalence of *T. gondii* in the prairie provinces of Canada, *T. gondii* is still present in the area. It is also possible that our negative findings are the result of infected individuals dying from the disease. Nonetheless, our results in conjunction with previous studies suggest that squirrels are not important intermediate hosts of *T. gondii* and likely do not act as reservoirs for *T. gondii* in cities. This knowledge is important as squirrels tend to occur at higher densities in cities than in more natural habitats which creates the possibility for increased parasite transmission between squirrels and cats, and further, cats and humans. When wildlife parasites are a human health concern, such as with *T. gondii*, then management actions are warranted. Our results suggest that squirrels do not need to be considered as sources of infection to cats. Consequently, no management efforts are needed in cities with abundant squirrel populations. In general, our results shed light on the prevalence of *T. gondii* in Sciurids in urban areas. This is important as *T. gondii* infection dynamics are still relatively unknown in cities.

## Acknowledgments

We want to thank Neil Pople, Amanda Salo, Niaz Rahim, and other members of the Manitoba Veterinary Diagnostic Services Laboratory for their helpful guidance and data collection. We thank Mitchell Green, Kyle Lefort, and Paul O’Brien for their help with necropsies, and Constance Finney and Emily Jenkins for their excellent suggestions regarding the methods. We want to give special thanks to Vera and Phil Froese for giving their permission to work on their land.

## References

1. Mackenstedt U, Jenkins D, Romig T. The role of wildlife in the transmission of parasitic zoonoses in peri-urban and urban areas. Int J Parasitol Parasites Wildl 2015;4:71–9. doi:10.1016/j.ijppaw.2015.01.006.

2. Deplazes P, Hegglin D, Gloor S, Romig T. Wilderness in the city: The urbanization of Echinococcus multilocularis. Trends Parasitol 2004;20:77–84. doi:10.1016/j.pt.2003.11.011.

3. Reperant LA, Hegglin D, Tanner I, Fischer C, Deplazes P. Rodents as shared indicators for zoonotic parasites of carnivores in urban environments. Parasitology 2009;136:329–37. doi:10.1017/S0031182008005428.

4. Lehrer EW, Fredebaugh SL, Schooley RL, Mateus-Pinilla NE. Prevalence of antibodies to Toxoplasma gondii in woodchucks across an urban-rural gradient. J Wildl Dis 2010;46:977–80. doi:10.7589/0090-3558-46.3.977.

5. Giraudeau M, Mousel M, Earl S, McGraw K. Parasites in the city: Degree of urbanization predicts poxvirus and coccidian infections in house finches (Haemorhous mexicanus). PLoS One 2014;9. doi:10.1371/journal.pone.0086747.

6. Gliwicz J, Goszczynski J, Luniak M. Characteristic features of animal populations under synurbanization - the case of the blackbird and of the striped field mouse. Memorab Zool 1994;49:237–44.

7. Bradley CA, Altizer S. Urbanization and the ecology of wildlife diseases. Trends Ecol Evol 2007;22:95–102. doi:10.1016/j.tree.2006.11.001.

8. Baker PJ, Molony SE, Stone E, Cuthill IC, Harris S. Cats about town: Is predation by free-ranging pet cats Felis catus likely to affect urban bird populations? Ibis (Lond 1859) 2008;150:86–99. doi:10.1111/j.1474-919X.2008.00836.x.

9. Sims V, Evans KL, Newson SE, Tratalos JA, Gaston KJ. Avian assemblage structure and domestic cat densities in urban environments. Divers Distrib 2008. doi:10.1111/j.1472-4642.2007.00444.x.

10. Elmore SA, Jones JL, Conrad PA, Patton S, Lindsay DS, Dubey JP. Toxoplasma gondii: epidemiology, feline clinical aspects, and prevention. Trends Parasitol 2010;26:190–6. doi:10.1016/j.pt.2010.01.009.

11. Dubey JP. Oocyst shedding by cats fed isolated bradyzoites and comparison of infectivity of bradyzoites of the VEG strain Toxoplasma gondii to cats and mice. J Parasitol 2001;87:215–9. doi:10.2307/3285204.

12. Fayyad A, Kummerfeld M, Davina I, Wohlsein P, Beineke A, Baumgärtner W, et al. Fatal systemic Toxoplasma gondii infection in a red squirrel (Sciurus vulgaris), a Swinhoe’s striped squirrel (Tamiops swinhoei) and a New World porcupine (Erethizontidae sp.). J Comp Pathol 2016;154:263–7. doi:10.1016/j.jcpa.2016.02.002.

13. Centers for Disease Control and Prevention. Toxoplasmosis 2019. Available from: https://www.cdc.gov/parasites/toxoplasmosis/.

14. Tenter AM, Heckeroth AR, Weiss LM. Toxoplasma gondii: From animals to humans. Int J Parasitol 2000;30:1217–58. doi:10.1016/S0020-7519(00)00124-7.

15. Dixon BR. Prevalence and control of toxoplasmosis - a Canadian perspective. Food Control 1992;3:68–75. doi:10.1016/0956-7135(92)90034-8.

16. Hill DE, Chirukandoth S, Dubey JP. Biology and epidemiology of Toxoplasma gondii in man and animals. Anim Heal Res Rev 2005;6:41–61. doi:10.1079/ahr2005100.

17. Conrad PA, Miller MA, Kreuder C, James ER, Mazet J, Dabritz H, et al. Transmission of Toxoplasma: Clues from the study of sea otters as sentinels of Toxoplasma gondii flow into the marine environment. Int J Parasitol 2005;35:1155–68. doi:10.1016/j.ijpara.2005.07.002.

18. Ballash GA, Dubey JP, Kwok OCHH, Shoben AB, Robison TL, Kraft TJ, et al. Seroprevalence of Toxoplasma gondii in White-tailed deer (Odocoileus virginianus) and free-roaming cats (Felis catus) across a suburban to urban gradient in northeastern Ohio. Ecohealth 2015;12:359–67. doi:10.1007/s10393-014-0975-2.

19. Parker TS, Nilon CH. Gray squirrel density, habitat suitability, and behavior in urban parks. Urban Ecosyst 2008. doi:10.1007/s11252-008-0060-0.

20. Dubey JP, Hodgin EC, Hamir AN. Acute Fatal Toxoplasmosis in Squirrels (Sciurus carolensis) with Bradyzoites in Visceral Tissues. J Parasitol 2006;92:658–9. doi:10.1645/GE-749R.1.

21. Jokelainen P, Nylund M. Acute fatal Toxoplasmosis in three Eurasian Red Squirrels (Sciurus vulgaris) caused by genotype II of Toxoplasma gondii. J Wildl Dis 2012;48:454–7. doi:10.7589/0090-3558-48.2.454.

22. Dubey JP, Frenkel JK. Experimental toxoplasma infection in mice with strains producing oocysts. J Parasitol 1973;59:505–12. doi:10.2307/3278784.

23. Afonso E, Thulliez P, Pontier D, Gilot-Fromont E. Toxoplasmosis in prey species and consequences for prevalence in feral cats: Not all prey species are equal. Parasitology 2007;134:1963–71. doi:10.1017/S0031182007003320.

24. Galeh TM, Sarvi S, Montazeri M, Moosazadeh M, Nakhaei M, Shariatzadeh SA, et al. Global status of Toxoplasma gondii seroprevalence in rodents: A systematic review and meta-analysis. Front Vet Sci 2020;7:461. doi:10.3389/fvets.2020.00461.

25. Statistics Canada. Winnipeg, CY, Manitoba. Census Profile. 2016 Census. Statistics Canada Catalogue no. 98-316-X2016001. 2016. Available from: https://www12.statcan.gc.ca.

26. Environment Canada. Canadian Climate Normals 1981–2010 Station Data for Winnipeg. 2020. Available from: https://climate.weather.gc.ca/climate_normals/results_1981_2010_e.html.

27. Kahle D, Wickham H. ggmap: Spatial visualization with ggplot2. R J 2013;5:144–61. doi:10.32614/rj-2013-014.

28. Elmore SA, Huyvaert KP, Bailey LL, Iqbal A, Su C, Dixon BR, et al. Multi-scale occupancy approach to estimate Toxoplasma gondii prevalence and detection probability in tissues: an application and guide for field sampling. Int J Parasitol 2016;46:563–70. doi:10.1016/j.ijpara.2016.04.003.

29. De Craeye S, Speybroeck N, Ajzenberg D, Dardé ML, Collinet F, Tavernier P, et al. Toxoplasma gondii and Neospora caninum in wildlife: Common parasites in Belgian foxes and Cervidae? Vet Parasitol 2011;178:64–9. doi:10.1016/j.vetpar.2010.12.016.

30. Tizard IR, Harmeson J, Lai CH. The prevalence of serum antibodies to Toxoplasma gondii in Ontario mammals. Can J Comp Med Rev Can Med Comp 1978;42:177–83.

31. Smith DD, Frenkel JK. Prevalence of antibodies to Toxoplasma gondii in wild mammals of Missouri and east central Kansas: biologic and ecologic considerations of transmission. J Wildl Dis 1995;31:15–21. doi:10.7589/0090-3558-31.1.15.

32. Lindsay DS, Weston JL, Little SE. Prevalence of antibodies to Neospora caninum and Toxoplasma gondii in gray foxes (Urocyon cinereoargenteus) from South Carolina. Vet Parasitol 2001;97:159–64. doi:10.1016/S0304-4017(01)00390-9.

33. Wanha K, Edelhofer R, Gabler-Eduardo C, Prosl H. Prevalence of antibodies against Neospora caninum and Toxoplasma gondii in dogs and foxes in Austria. Vet Parasitol 2005;128:189–93. doi:10.1016/j.vetpar.2004.11.027.

34. Frenkel JK, Hassanein KM, Hassanein RS, Brown E, Thulliez P, Quintero-Nunez R. Transmission of Toxoplasma gondii in Panama City, Panama: A five-year prospective cohort study of children, cats, rodents, birds, and soil. Am J Trop Med Hyg 1995;53:458–68. doi:10.4269/ajtmh.1995.53.458.

35. Dubey JP, Dennis PM, Verma SK, Choudhary S, Ferreira LR, Oliveira S, et al. Epidemiology of toxoplasmosis in white tailed deer (Odocoileus virginianus): Occurrence, congenital transmission, correlates of infection, isolation, and genetic characterization of Toxoplasma gondii. Vet Parasitol 2014;202:270–5. doi:10.1016/j.vetpar.2014.01.006.

36. Mercier A, Garba M, Bonnabau H, Kane M, Rossi JP, Dardé ML, et al. Toxoplasmosis seroprevalence in urban rodents: A survey in Niamey, Niger. Mem Inst Oswaldo Cruz 2013;108:399–407. doi:10.1590/S0074-0276108042013002.

37. Murphy RG, Williams RH, Hughes JM, Hide G, Ford NJ, Oldbury DJ. The urban house mouse (Mus domesticus) as a reservoir of infection for the human parasite Toxoplasma gondii: An unrecognised public health issue? Int J Environ Health Res 2008;18:177–85. doi:10.1080/09603120701540856.

38. Jacobs L, Stanley AM, Herman CM. Prevalence of Toxoplasma antibodies in rabbits, squirrels, and raccoons collected in and near the Patuxent Wildlife Research Center. J Parasitol 1962;48:550. doi:10.2307/3274906.

39. Walton BC, Walls KW. Prevalence of toxoplasmosis in wild animals from Fort Stewart, Georgia, as indicated by serological tests and mouse inoculation. Am J Trop Med Hyg 1964;13:530–3. doi:10.4269/ajtmh.1964.13.530.

40. Smith DD, Frenkel JK. Prevalence of Antibodies To Toxoplasma Gondii in Wild Mammals of Missouri and East Central Kansas: Biologic and Ecologic Considerations of Transmission. J Wildl Dis 1995;31:15–21. doi:10.7589/0090-3558-31.1.15.

41. Soave OA, Lennette EH. Naturally acquired toxoplasmosis in the gray squirrel, sciurus griseus, and its bearing on the laboratory diagnosis of rabies. J Lab Clin Med 1959;53:163–6. doi:10.5555/uri:pii:0022214359900642.

42. Burridge MJ, Bigler WJ, Forrester DJ, Hennemann JM. Serologic survey for Toxoplasma gondii in wild animals in Florida. J Am Vet Med Assoc 1979;175:964–7.

43. Opsteegh M, Langelaar M, Sprong H, den Hartog L, De Craeye S, Bokken G, et al. Direct detection and genotyping of Toxoplasma gondii in meat samples using magnetic capture and PCR. Int J Food Microbiol 2010;139:193–201. doi:10.1016/j.ijfoodmicro.2010.02.027.

44. Gilbert AT, Fooks AR, Hayman DTS, Horton DL, Müller T, Plowright R, et al. Deciphering serology to understand the ecology of infectious diseases in wildlife. Ecohealth 2013;10:298–313. doi:10.1007/s10393-013-0856-0.

45. Roqueplo C, Halos L, Cabre O, Davoust B. Toxoplasma gondii in wild and domestic animals from New Caledonia. Parasite 2011;18:345–8. doi:10.1051/parasite/2011184345.

46. Sharma R, Parker S, Elkin B, Mulders R, Branigan M, Pongracz J, et al. Risk factors and prevalence of antibodies for Toxoplasma gondii in diaphragmatic fluid in wolverines (Gulo gulo) from the Northwest Territories, Canada. Food Waterborne Parasitol 2019;15:e00056. doi:10.1016/j.fawpar.2019.e00056.

47. Sharma R, Parker S, Al-Adhami B, Bachand N, Jenkins E. Comparison of tissues (heart vs. brain) and serological tests (MAT, ELISA and IFAT) for detection of Toxoplasma gondii in naturally infected wolverines (Gulo gulo) from the Yukon, Canada. Food Waterborne Parasitol 2019;15:e00046. doi:10.1016/j.fawpar.2019.e00046.

48. Hirota J, Nishi H, Matsuda H, Tsunemitsu H, Shimizu S. Cross-reactivity of Japanese encephalitis virus-vaccinated horse sera in serodiagnosis of West Nile Virus. J Vet Med Sci 2010;72:369–72. doi:10.1292/jvms.09-0311.

49. Sekla, L., Stackiw W, Rodgers S. A serosurvey of toxoplas-mosis in Manitoba. Can J Public Heal 1981;72:111–7.

50. Shettigara PT, Choi NW, Abu-Zeid HAH. Prevalence of toxoplasmosis in Manitoba province: a seroepidemiologic study of 23 146 prenatal sera. Am J Epidemiol 1976;104:340–340.

51. Fayer R. Toxoplasmosis update and public health implications. Can Vet J 1981;22:344–52.

52. Poljak Z, Dewey CE, Friendship RM, Martin SW, Christensen J, Ojkic D, et al. Pig and herd level prevalence of Toxoplasma gondii in Ontario finisher pigs in 2001, 2003, and 2004. Can J Vet Res 2008;72:303–10.

53. Nation PN, Allen JR. Antibodies to Toxoplasma gondii in Saskatchewan cats, sheep and cattle. Can Vet J 1976;17:308–10.

54. Smith HJ. Seroprevalence of anti-Toxoplasma IgG in Canadian swine. Can J Vet Res 1991;55:380–1.

55. Environment Canada. Monthly Data Report for 2007 for Winnipeg The Forks, Manitoba. 2020. Available from: https://climate.weather.gc.ca/climate_data/monthly_data_e.html.

56. Dubey JP, Miller NL, Frenkel JK. The toxoplasma gondii oocyst from cat feces. J Exp Med 1970;132:636–62. doi:10.1084/jem.132.4.636.

57. Frenkel JK, Ruiz A, Chinchilla M. Soil survival of Toxoplasma oocysts in Kansas and Costa Rica. Am J Trop Med Hyg 1975;24:439–43. doi:10.4269/ajtmh.1975.24.439.

58. Government of Manitoba. Manitoba Drought Management Strategy. 2020. Available from: https://www.gov.mb.ca/sd/water/drought_condition/index.html.

59. Lélu M, Villena I, Dardé ML, Aubert D, Geers R, Dupuis E, et al. Quantitative estimation of the viability of Toxoplasma gondii oocysts in soil. Appl Environ Microbiol 2012;78:5127–32. doi:10.1128/AEM.00246-12.

60. Hwang YT, Pitt JA, Quirk TW, Dubey JP. Seroprevalence of Toxoplasma gondii in mesocarnivores of the Canadian prairies. J Parasitol 2007;93:1370–3. doi:10.1645/GE-1319.1.

