## Supporting Information for "No evidence of *Toxoplasma gondii* infection in urban and rural squirrels"

*Authors:* Riikka P Kinnunen (RPK), Chloé Schmidt (CS), Adrián Hernández-Ortiz (AHO), Colin J Garroway (CJG)

*Affiliations:* ^1^Department of Biological Sciences, Biological Sciences Building, University of Manitoba, Winnipeg, MB, Canada R3T 2N2, ^2^ Department of Veterinary Microbiology, Western College of Veterinary Medicine, University of Saskatchewan, 52 Campus Drive, Saskatoon, SK, Canada S7N 5B4

Table S1. Primers and probe sequences used in the PCR to detect *Toxoplasma gondii* and cellular r18S DNA.

|  | Target | Name | Sequence 5_→ 3_ | 5_ Modification | 3_ Modification |
| --- | --- | --- | --- | --- | --- |
| Primers | *T. gondii* | T2 | CGGAGAGGGAGAAGATGTT |  |  |
|  | *T. gondii* | T3 | GCCATCACCACGAGGAAA |  |  |
|  | Cell r18S | F | GATTAAGTCCCTGCCCTTT |  |  |
|  | Cell r18S | R | GATAGTCAAGTTCGACCGTCTT |  |  |
| Probes | *T. gondii* |  | CTTGGCTGCTTTTCCTGGAGGG | FAM | BHQ1 |
|  | Cell r18S |  | CACACCGCCCGTCGCTACTACC | Cy5 | BHQ2 |

Table S2. Results of the enzyme-linked immunosorbent assay (ELISA) test carried out on the 15 serum samples from grey squirrels (*Sciurus carolinensis*) to detect IgG serum antibodies to *Toxoplasma gondii.* Sample to Positive Ratio (S/P) percentage (S/P%) was calculated for each sample. Samples with S/P% less or equal to 40% were considered negative; samples with S/P% between 40 and 50% doubtful or inconclusive; and samples with an S/P% higher than 50% positive.

| Number | ID | S/P % | Result |
| --- | --- | --- | --- |
| Sq01 | 268-19 | 0.7 | N |
| Sq02 | 327-19 | -0.1 | N |
| Sq03 | 326-19 | -1 | N |
| Sq04 | 329-19 | -0.6 | N |
| Sq05 | 342-19 | 0.4 | N |
| Sq06* | 343-19 | 0.5 | N |
| Sq07 | 344-19 | -1 | N |
| Sq08 | 347-19A | -0.4 | N |
| Sq09 | 347-19B | -0.6 | N |
| Sq10 | 351-19 | -0.6 | N |
| Sq11 | 355-19 | -1.1 | N |
| Sq12 | 359-19 | -1.1 | N |
| Sq13 | 389-19 | 0.8 | N |
| Sq14* | 391-19 | -0.7 | N |
| Sq15 | 392-19 | -0.5 | N |

* Not enough sample
